# Over-expression of Cyclic Nucleotide-Gated Ion Channel 2 (CNGC2) triggers hypersensitivity to virulent pathogens and elevated Ca^2+^

**DOI:** 10.1101/2025.10.29.685368

**Authors:** Sonhita Chakraborty, Hyunsuh Lee, Eiji Nambara, Wolfgang Moeder, Keiko Yoshioka

## Abstract

The Arabidopsis *Cyclic Nucleotide-Gated Ion Channel 2* (CNGC2), also known as *Defense No Death 1* (DND1), is the most extensively studied plant CNGC and has been implicated in diverse physiological processes, including floral transition, responses to heat and humidity, and hormone signaling. Its role in immunity has received particular attention due to the autoimmunity phenotype observed in the *cngc2/dnd1* knockout mutants. Interestingly, despite this hyperactivation of immunity, the mutant also exhibits impaired hypersensitive cell death—a hallmark of effector-triggered immunity (ETI)—as well as reduced reactive oxygen species (ROS) production and diminished Ca²⁺ influx in response to pathogen-associated molecular patterns (PAMPs) such as the bacterial flagellin peptide flg22. These contradictory phenotypes highlight the complex biological functions of CNGC2.

To date, most studies have focused on loss-of-function mutants. In this study, we performed a detailed characterization of CNGC2 overexpression lines to gain deeper insight into its role in immunity. Remarkably, overexpression of CNGC2 led to heightened susceptibility to two taxonomically distinct pathogens, despite the plants displaying wild-type morphology. Overexpression of CNGC2 rescued several *cngc2* mutant phenotypes, including morphological defects and delayed flowering, yet these plants were also hypersensitive to elevated external Ca²⁺ levels. Furthermore, they exhibited attenuated responses to flg22, suggesting that CNGC2 does not act as a simple positive or negative regulator of immunity. Our findings reveal an essential role for CNGC2 where a balanced expression level is critical for maintaining Ca²⁺ homeostasis between the apoplast and cytosol, thereby influencing the generation of Ca²⁺ signals essential for immune responses.

## Introduction

Like all eukaryotes, plants utilize Calcium ions (Ca^2+^) as a second messenger for many signaling pathways. This is achieved by a Ca^2+^ concentration gradient between the apoplastic space and cytosol: the cytosolic Ca^2+^ concentration [Ca^2+^]_cyt_ is maintained 10,000-fold lower than that in the apoplast (Demidchik et al., 2018). Thus, the opening of Ca^2+^ channels leads to a rapid Ca^2+^ influx and an increase in [Ca^2+^]_cyt_, which acts as a Ca^2+^ signal. Ca^2+^ is then removed from the cytosol by Ca^2+^ pumps and transporter proteins, like vacuolar H^+^ /Cation exchangers (CAX), to go back to the resting state and the cell is ready for another stimulus (Demidchik et al., 2018). It is believed that different stimuli generate distinct spatio-temporal patterns of Ca^2+^ influx, also called a Ca^2+^ signature, which is then sensed by calcium sensor proteins such as calmodulins (CaM), calcium-dependent protein kinases (CDPKs) and others, which transduce the signal into specific downstream responses (DeFalco et al., 2010). In recent years, the role of Ca^2+^ signaling in immunity has seen significant progress and a number of conventional and non-conventional Ca^2+^ channels have been connected to immune responses (Xu et al., 2022; Li et al., 2024).

In plant immunity, the first layer of defense is through the recognition of common microbial molecules, such as bacterial flagellin or fungal chitin, termed pathogen-associated molecular patterns (PAMPs). This occurs by plasma membrane (PM) localized pattern recognition receptors (PRR), which trigger PAMP-triggered immunity (PTI). The best studied example is the recognition of a 22 amino acid epitope of the bacterial flagellin (flg22) by the PRR, FLS2 (DeFalco and Zipfel, 2021). To overcome PTI, many virulent pathogens have evolved effector proteins that, can dampen the plant immune response; in turn, plants have evolved a second layer of defense, where these effector proteins are recognized by cytosolic NLR (nucleotide binding-leucine rich repeat) receptor proteins, which triggers a stronger type of immune response called effector induced immunity (ETI) (Weralupitiya et al., 2024). Recent data suggests that both PTI and ETI trigger somewhat overlapping responses, including a rapid Ca^2+^ influx and the production of reactive oxygen species (ROS) (Ngou et al., 2021; Xu et al., 2022). Members of the Cyclic Nucleotide-Gated Ion Channel (CNGC), osmotic stress-induced calcium (OSCA) and Glutamate-like receptor (GLR) families have been connected to PTI (Tian et al., 2019; Thor et al., 2020; Bjornson et al., 2021), while recent data suggest that during ETI some NLRs oligomerize to form multimeric resistosomes that facilitate a prolonged Ca^2+^ influx triggering programmed cell death (PCD) in the Hypersensitive Response (HR) (Kim et al., 2022; Weralupitiya et al., 2024).

Cyclic Nucleotide-Gated Ion Channels (CNGC) are among the best studied Ca^2+^ channels (Dietrich et al., 2020; Jarratt-Barnham et al., 2021). They have been implicated in a variety of physiological processes, such as pollen tube and root tip growth, thermo- and humidity-sensing, symbiotic and pathogenic plant microbe interactions (Charpentier et al., 2006; Finka et al., 2012; Brost et al., 2019; Tian et al., 2019; Tan et al., 2020; Hussain et al., 2024). Plant CNGCs – like their animal counterparts – are believed to form tetrameric channels, which could be comprised of either one type (homomeric) or different subunits (heteromeric). The latter has been shown for CNGCs 2 and 4 (Chin et al., 2013; Tian et al., 2019) and CNGCs 18 and 8 (Pan et al., 2019). CNGC activity is regulated by the binding of CaMs, which can have both positive or negative effects on channel activity (DeFalco et al., 2016; Pan et al., 2019; Tian et al., 2019) or via phosphorylation (Tian et al., 2019; Sun et al., 2025; Zhu et al., 2025; Yang et al.).

The best-studied CNGCs are the two closely related Arabidopsis CNGCs, CNGC2 and CNGC4. Their null mutants were initially identified as *defense no death* (*dnd1*) and *dnd2* (also named *HR-like lesion mimic* (*hlm1*), (Clough et al., 2000; Balagué et al., 2003; Jurkowski et al., 2004). They show a reduced HR response when infected with pathogens that trigger ETI. Both mutants also display similar autoimmune phenotypes, including conditional spontaneous cell death, increased accumulation of the defense hormone salicylic acid (SA), and enhanced resistance against biotrophic and necrotrophic pathogens. Both mutants exhibit the constitutive activation of SA and jasmonic acid (JA) pathways (Clough et al., 2000; Jurkowski et al., 2004; Genger et al., 2008).

Another CNGC-related mutant, *constitutive expresser of pathogenesis-related genes 22* (*cpr22*), which is a gain-of-function mutant of CNGC11 and 12, also displays autoimmunity phenotypes (Yoshioka et al., 2006). Its autoimmunity phenotypes are caused by the elevated [Ca^2+^]_cyt_ levels and can be supressed by Ca^2+^ channel blockers, indicating the constitutive Ca^2+^ influx in *cpr22* hyper-activates immunity (Urquhart et al., 2007; Moeder et al., 2019). In contrast, *cngc2* displays an impaired HR phenotype and reduced Ca^2+^ influx upon PAMP treatment, which gave rise to the notion that CNGC2 is a positive regulator of defense (Ali et al., 2007; Tian et al., 2019). However, this defect in PTI is only seen when plants are grown on media at standard Ca^2+^ concentration (1.5 mM), while plants grown at very low (0.1 mM) external Ca^2+^ do not display this phenotype (Wang et al., 2017; Tian et al., 2019). On the other hand, the autoimmunity phenotype of *cngc2*, which is a loss of function mutant, rather suggests a role as a negative regulator (Moeder et al., 2011). This is also supported by several publications showing that *CNGC2* is transcriptionally downregulated when immunity is activated. Zhu et al. (2010) show that the transcriptional corepressor Topless-related 1 (TPR1) represses the expression of *CNGC2* (and *CNGC4*) during pathogen infection. (Niu et al., 2019) further showed that exogenous SA treatment also reduces *CNGC2* (and *CNGC4*) expression.

Interestingly, *cngc2* knockout mutants display a number of phenotypes that are not typical for autoimmune mutants, such as hyper-susceptibility to elevated Ca^2+^ (Chan et al., 2003; Chan et al., 2008) and delayed flowering (Chin et al., 2013; Fortuna et al., 2015). It was also shown that CNGC is part of a negative feedback loop that regulates auxin homeostasis (Chakraborty et al., 2021). Recent data shows that CNGC2 plays crucial roles in development (Wang et al., 2022b), responses to DAMPs (Wang et al., 2022a), and environmental changes (Finka et al., 2012; Hussain et al., 2024), suggesting CNGC2 has a broader role in Ca²⁺ homeostasis and may act as a nexus integrating diverse stimuli in plants.

Thus, in this study, we over-expressed *CNGC2* (via the CaMV35S promoter) in *cngc2* plants and found that, unexpectedly, these plants were hyper-susceptible to the two different virulent pathogens *Hyaloperonospora arabidopsidis* Noco2 and *Pseudomonas syringae* <u>pv.</u> DC3000. The overexpression complemented some of the *cngc2* phenotypes, like the autoimmunity and delayed flowering, while these plants were also hyper-susceptible to elevated Ca^2+^ in the medium and displayed elevated endogenous IAA like *cngc2*, suggesting a more complex role for CNGC2 in Ca^2+^ homeostasis.

## Results

### Ectopic overexpression of CNGC2 reverts pleiotropic phenotypes of *cngc2*

*cngc2/dnd1* plants display an impaired HR phenotype and reduced Ca^2+^ influx upon PAMP treatment (Tian et al., 2019). However, the fact that a loss-of-function mutant exhibits autoimmunity (Yu et al., 1998) as well as the observed downregulation of *CNGC2* after pathogen infection (Zhu et al., 2010; Moeder et al., 2011; Niu et al., 2019) or flg22 treatment suggests CNGC2 is rather a negative regulator of immunity (Fig.1). To address this question, we introduced *CNGC2* under the control of the constitutive CaMV 35S promoter into the *cngc2* mutant background (*cngc2 pCaMV 35S::CNGC2-YFP,* hereafter *CNGC2OX*). Note: We have previously shown that the presence of YFP at the C-terminus does not interfere with CNGC function (Yoshioka et al., 2006; Wang et al., 2022a). We analyzed three independent homozygous *CNGC2OX* lines. All three lines displayed elevated *CNGC2* expression compared to Col wild type (Fig. 2a). Further, confocal microscopy confirmed that CNGC2 protein is expressed and localizes to the plasma membrane (Fig. 2b). All three *CNGC2OX* lines rescued the *cngc2* morphological defects such as dwarfisms and abnormal leaf shape and were morphologically indiscernible from wild type plants (Fig. 3A top). The delayed flowering transition seen in *cngc2* was also rescued in *CNGC2OX* lines (Fig. 3B). These data indicated that all developmental defects of *cngc2* mutant were rescued by over expression of *CNGC2*. The autoimmunity-related spontaneous cell death and elevated basal levels of endogenous SA observed in *cngc2* plants were also suppressed in the *CNGC2OX* lines and were comparable to those of Col wild type plants (Fig. 3A bottom, 3C). It is well documented that in the presence of the bacterial PMP flg22, Col wild type plants display an inhibition of root growth (Chinchilla et al., 2007). We found that *cngc2* seedlings grown on MS plates supplemented with 1 µM flg22 were insensitive against flg22, while wild type plants showed stunted roots as expected. All three *CNGC2OX* lines displayed wild type-like flg22 sensitivity (Fig. 3D). All these results suggest that the overexpression of CNGC2 complemented *cngc2* developmental and autoimmune phenotypes.

**Fig 1.**
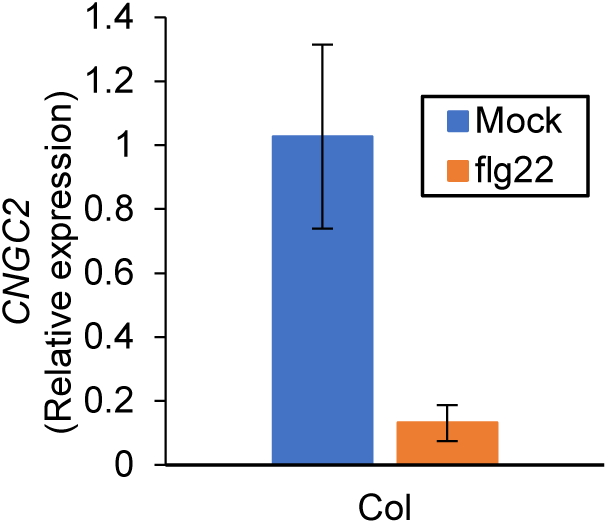
*CNGC2* is transcriptionally down-regulated upon flg22 treatment. Q-PCR analysis of *CNGC2* expression in 6-week-old Col wild type plants after treatment with 1 µM flg22. Transcripts were normalized to *AtEF1A*. Each bar represents the mean of three biological repeats ± SE.

**Figure 2.**
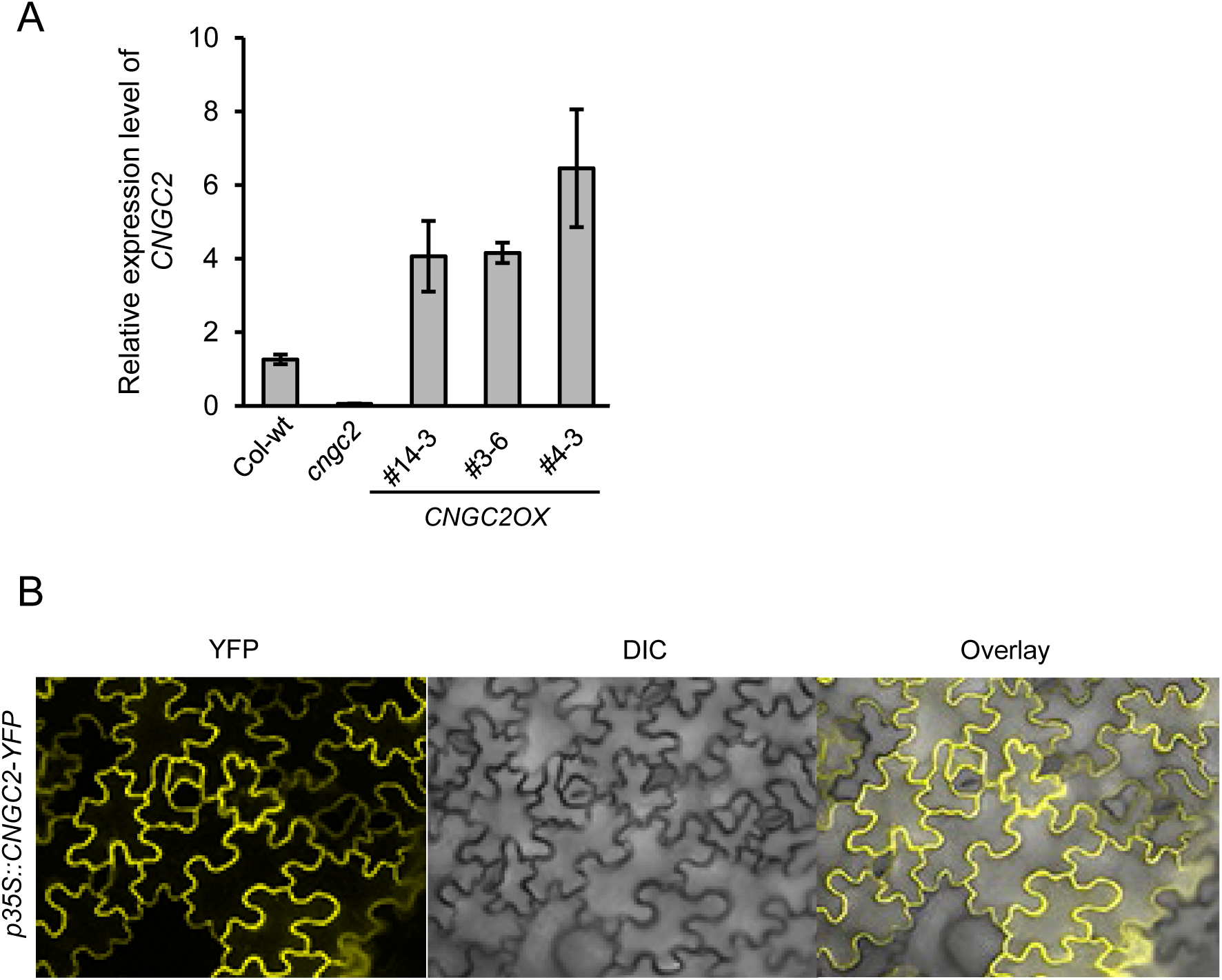
*CNGC2OX* lines express elevated levels of CNGC2. (A) CNGC2 expression levels in 3- to 4-week-old wild-type and *cngc2 35S::CNGC2-YFP* (*CNGC2OX)* leaves. (B) *Agrobacterium*-mediated transient expression of CNGC2-YFP in *Nicotiana benthamiana* shows subcellular localization of CNGC2-YFP to the plasma membrane.

**Figure 3.**
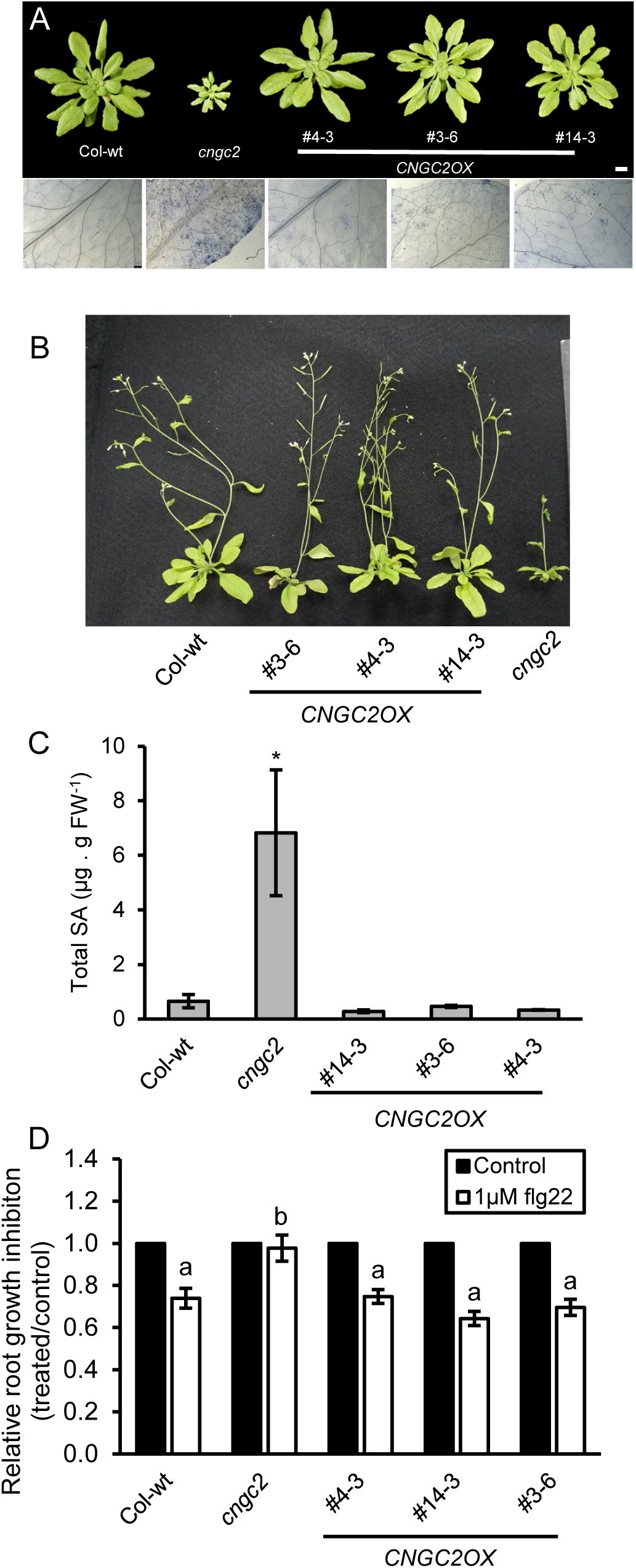
Ectopic over expression of CNGC2 rescues developmental phenotype. (A) (top) Morphology of *35S::CNGC2:YFP cngc2 (CNGC2OX)* lines, compared to Col-wt and *cngc2* when grown in short day conditions. Scale = 1cm. (bottom) Trypan blue staining of *CNGC2OX,* Col-wt *and cngc2* plants. Scale = 1mm (B) Flowering transition time of *CNGC2OX,* Col-wt *and cngc2* plants. (C) Total salicylic acid (SA) levels in 3- to 4-week-old Col-wt, *cngc2,* and *CNGC2OX* plants. Error bars indicate SE of three replicates. Bars marked with asterisks indicate significant differences (Student’s t test, P<0.05). (D) flg22-mediated root growth inhibition. Col-wt, *cngc2* and *CNGC2OX* seedlings were grown on ½ MS media (control) and media supplemented with 1uM flg22. Total root length was measured 12 days post germination. *cngc2* seedlings displayed reduced sensitivity in root growth compared to Col-wt. Root inhibition of *CNGC2OX* seedlings were similar to Col-wt. Error bars indicate SE of three replicates. Bars marked with different letters indicate significant differences (Tukey’s HSD test and pairwise comparison, p < 0.05).

### Overexpression of CNGC2 causes hyper-susceptibility to virulent pathogens and a diminished PTI

Despite their impaired PTI response and their HR-deficient phenotype, *cngc2* mutant plants exhibit enhanced resistance to virulent strains of the bacterial pathogen *Pseudomonas syringae* pv. DC3000 (*Pst* DC3000) and the oomycete *H. arabidopsidis* (Hpa; Genger et al., 2008; Clough et al., 2000). Since the morphological phenotypes and elevated SA levels associated with autoimmunity were reverted in the *CNGC2OX* lines, we tested their pathogen infection phenotypes against *Pst* DC3000. The autoimmune *cngc2* plants exhibited reduced growth of *Pst* as expected. However, while CNGC2 had been reported as a positive regulator of immunity, the *CNGC2OX* lines allowed even higher bacterial growth than wild type plants (Fig 4A), suggesting that overexpression of CNGC2 leads to hyper-susceptibility to virulent pathogens. This was also confirmed using the oomycete pathogen *Hpa* Noco2, which displayed much more hyphal growth and higher levels of sporangiophore formation than Col wild type (Fig. 4B, C). If CNGC2 were indeed a positive regulator of pathogen resistance, then we would expect that the *CNGC2OX* lines display enhanced levels of pathogen resistance; however, our data shows the opposite, arguing for a role as a negative regulator of immunity.

**Figure 4.**
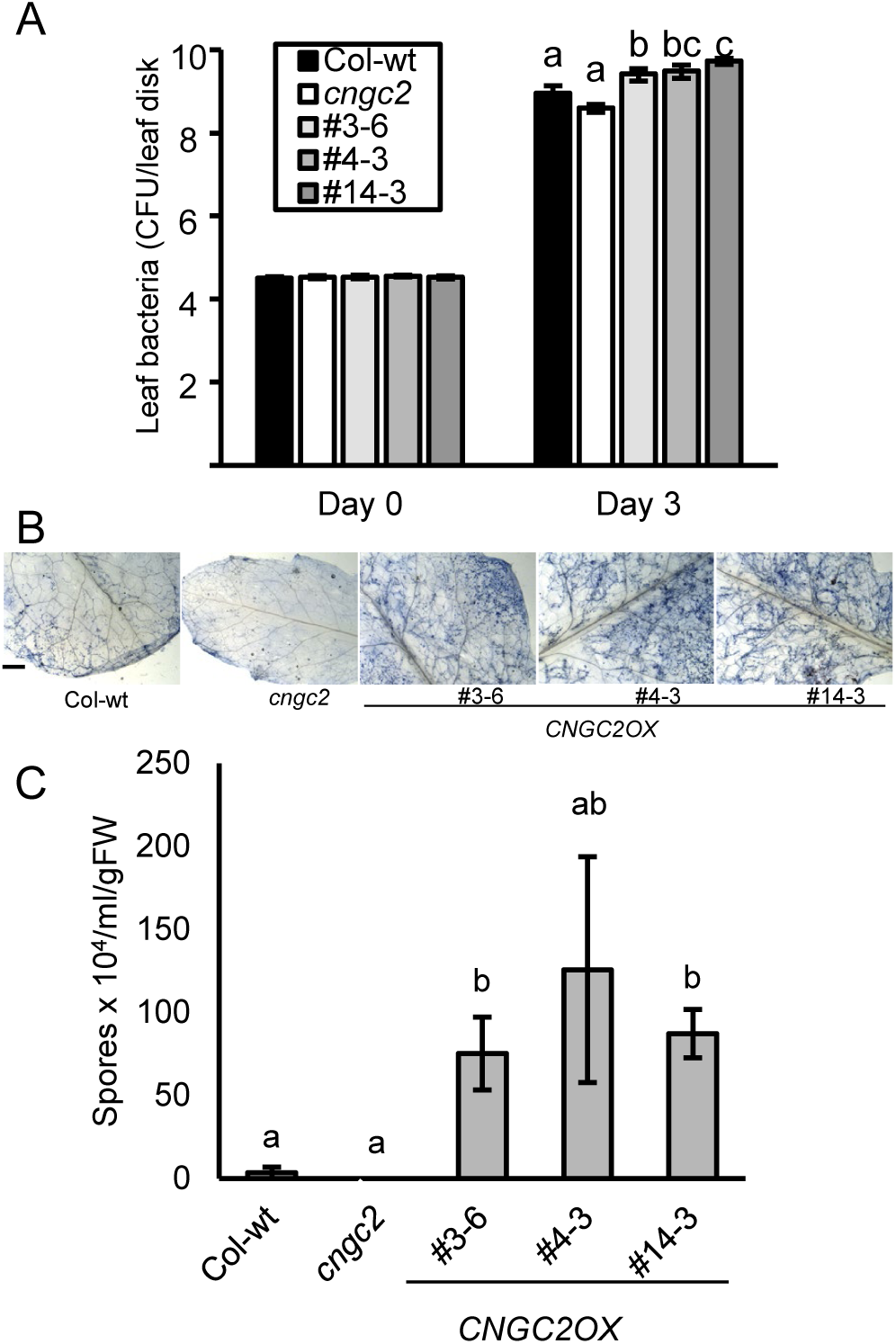
Ectopic over expression of CNGC2 induces enhanced pathogen susceptibility against virulent pathogens. (A) Leaf bacterial populations were assessed 3 d after infiltration of *CNGC2OX*, Col-wt and *cngc2* plants with virulent *P. syringae* pv. *tomat*o DC3000 at 5 × 10^4^ colony-forming units (cfu) mL^−1^. (B) Trypan blue staining of *plants infected with Hyaloperonospora arabidopsidis*, isolate Noco2, reveals increased sporangiophore formation and hyphae growth in the *CNGC2OX* lines. Scale = 1mm. (C) Spore count of *H. arabidopsidis* infected plants. Error bars indicate SE of three replicates. Bars marked with different letters indicate significant differences (Tukey’s HSD test and pairwise comparison, p < 0.05).

As previous data had shown that functional CNGC2 is required for a successful PTI response, evidenced by reduced increases in [Ca^2+^]_cyt_ and ROS burst after flg22 treatment (Tian et al., 2019), we next assessed these parameters in our *CNGC2OX* lines. As expected, *cngc2* mutant plants exhibited an attenuated ROS burst in response to flg22 treatment. However, surprisingly, the *CNGC2OX* lines also displayed a reduced ROS production (Fig. 5A). This observation led us to test their Ca^2+^ signal upon flg22 treatment. The *CNGC2OX* lines showed a significantly suppressed Ca^2+^ signal upon flg22 treatment, which was similar to *cngc2* (Fig. 5B). This data clearly shows that, surprisingly, elevated levels of this Ca^2+^ channel led to a reduction in Ca^2+^ influx upon PAMP treatment.

**Figure 5.**
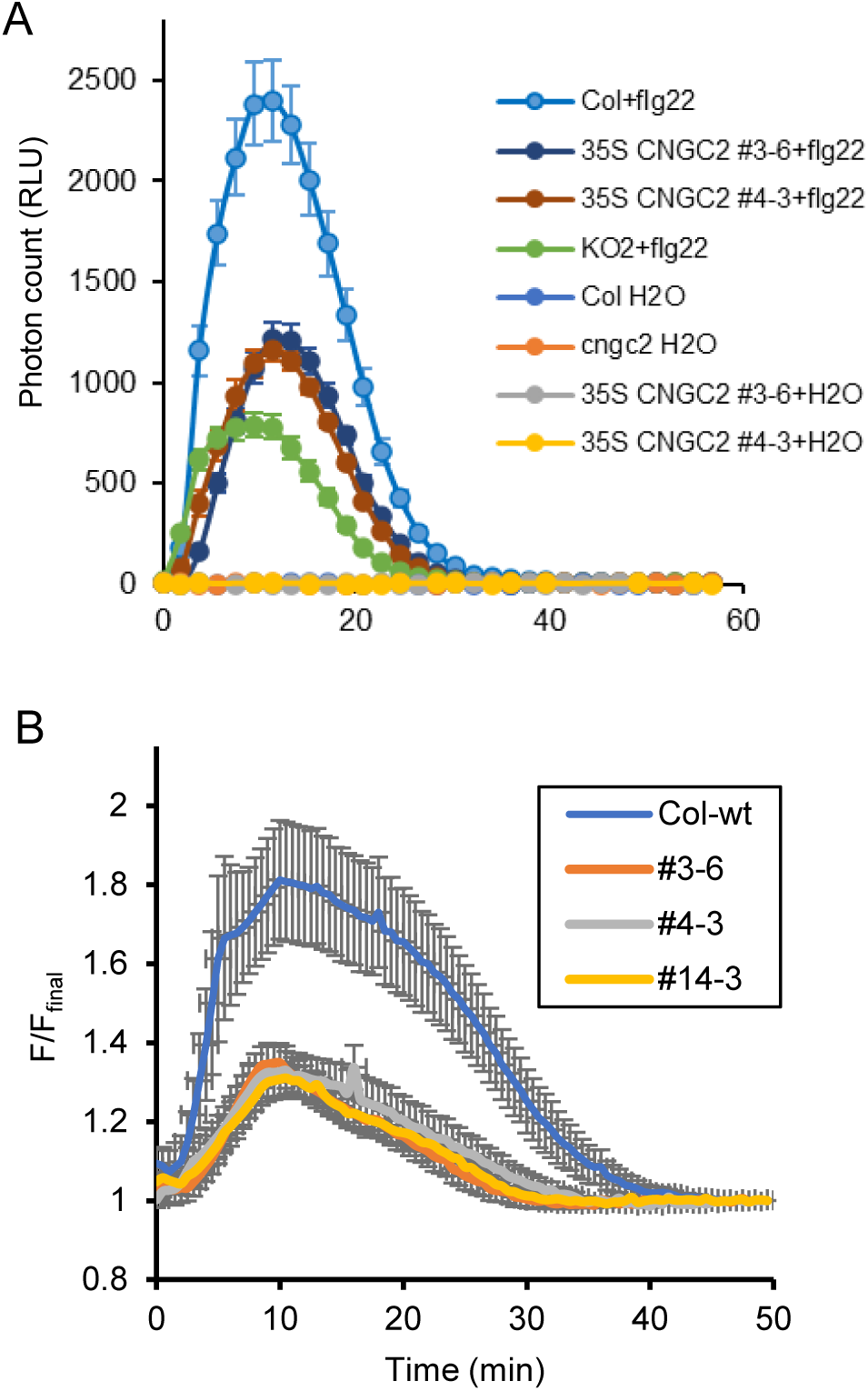
CNGC2OX plants exhibit an impaired PTI response. (A) flg22-trigged ROS production upon application of 100nM flg22 to leaf discs of Col-wt, *cngc2* and *CNGC2OX* plants Shown is the average ± SE of 20 replicates (B) Ca^2+^ signal upon application of 100nM flg22 to leaf discs of Col-wt, *cngc2* and *CNGC2OX* plants carrying the UBQ::GCaMP3 reporter. Shown is the average ± SE of 6 replicates.

### *CNGC2OX* plants are hypersensitive to elevated external Ca^2+^

*cngc2* mutants have been known for a long time to be hypersensitive to elevated Ca^2+^ in the medium (Chan et al., 2003; Chan et al., 2008). *cngc2* (and *cngc4*) knockout mutants are the only CNGC knockout mutants that display this Ca^2+^ sensitivity phenotype (Wang et al., 2017). Interestingly, the *repressor of defense, no death1* (*rdd1,* Chin et al. 2013) mutation, which alleviated most *cngc2* phenotypes like increased pathogen resistance and delayed floral transition, had no effect on *cngc2*’s Ca^2+^ sensitivity (Chin et al., 2013). When we tested our *CNGC2OX* lines on 0.5 x MS media plates supplemented with 20 mM CaCl_2_, we found that they displayed an even greater degree of Ca^2+^ sensitivity than *cngc2* plants (Fig. 6). This suggests that CNGC2 plays a role in Ca^2+^ homeostasis, and a reduction as well as an increase in CNGC2 channel protein has negative effects on the plant.

**Figure 6.**
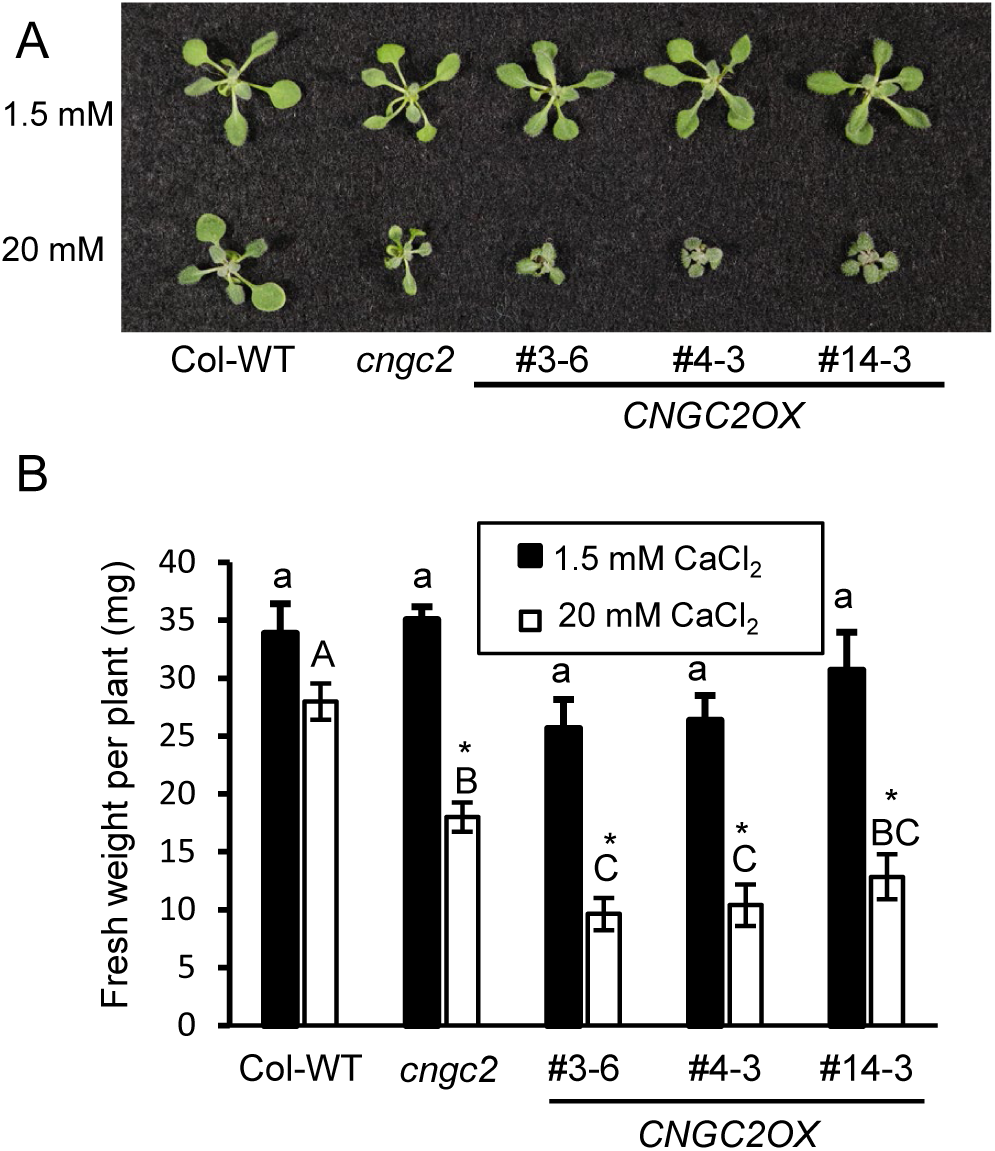
*CNGC2OX* plants are hypersusceptible to elevated external Ca^2+^. (A, B) Col wild-type (Col-wt), cngc2 and *CNGC2OX* plants were grown on both control (1.5 mM CaCl_2_) and medium supplemented with 20 mM CaCl_2_. (A) At 15 days photos were taken and (B) Shoots were weighed. Shown is the average weight/plant. Error bars indicate SE of four replicate plates. Asterisks indicate significance within each genotype (t-test (p<0.05). Different letters indicate significant differences within each treatment (ANOVA followed by Tukey’s HSD).

It has been reported that *cngc2* plants show elevated auxin levels of the endogenous auxin indole-3-acetic acid (IAA) (Chakraborty et al., 2021). Furthermore, the *rdd1* mutation, which causes a loss-of-function of the auxin biosynthesis gene *YUCCA6* (*YUC6*), partially suppressed many *cngc2* phenotypes. In the *rdd1 cngc2* double mutant, not only were the elevated IAA levels reverted to wild type levels, but a partial rescue of the *cngc2* autoimmunity phenotypes was also observed (Chin et al., 2013), suggesting that the elevated IAA levels in *cngc2* contribute to its pleotropic phenotypes. Thus, we also measured the IAA content of two *CNGC2OX* lines and found that they also displayed elevated IAA levels, similarly to *cngc2* (Fig. 7). This result further supports the notion that CNGC2 is not a simple positive regulator of immune responses, but rather is part of a homeostasis mechanism that affects immunity as well as balancing cellular auxin perception and biosynthesis.

**Figure 7.**
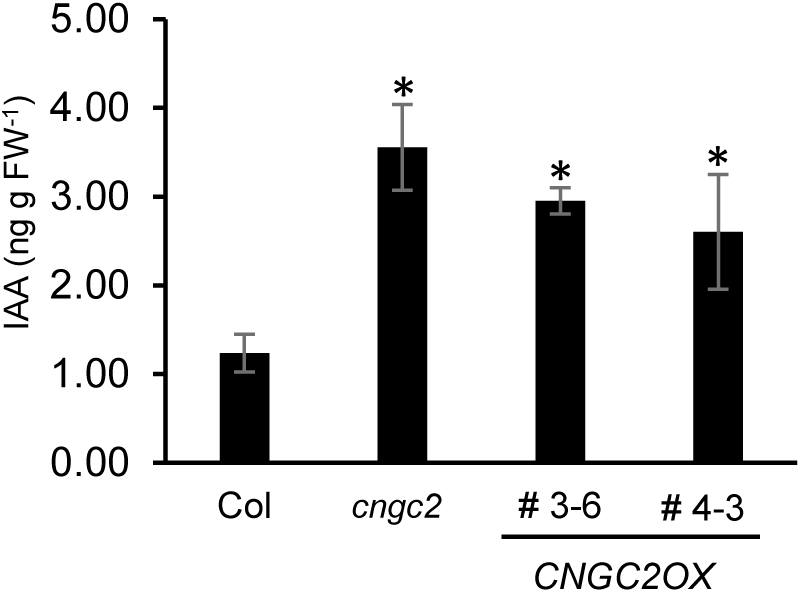
*CNGC2OX* plants exhibit elevated IAA levels. Shoot IAA levels of 5-week-old plants were measured using LC-MS/MS. The experiment was repeated two times and averages from one representative trial are presented; shown are means ± SE, n 3-4. Bars marked with an asterisk indicate significant difference from Col-wt (Student’s t test, P < 0.05).

## Discussion

Because of its toxic effects, cells keep the level of Ca^2+^ in the cytosol very low (10^−7^ M) and maintain a steep concentration gradient (10,000-fold). Cells then started to use Ca^2+^ as a second messenger, where the influx of Ca^2+^ through Ca^2+^ channels triggers physiological responses to many stimuli. The incoming Ca^2+^ is then rapidly removed from the cytosol creating the resting state again (Demidchik et al., 2018). This is accomplished by the coordinated action of influx Ca^2+^ channels and transporters that export Ca^2+^ against the gradient out of the cell. In animals, the endoplasmic reticulum and mitochondria serve as Ca^2+^ repositories, while plants mostly utilize the apoplast and vacuole for Ca^2+^ sequestration (DeFalco et al., 2010).

The influx of Ca^2+^ immediately triggers the mechanisms to remove it from the cytosol since prolonged elevation of [Ca^2+^]_cyt_ has detrimental effects. This can be seen in plants that express constitutive active mutants of Ca^2+^ channels, like *cpr22* (a mis-regulated fusion of CNGC11 and 12) (Yoshioka et al., 2006) and *cngc20-4* (Zhao et al., 2021) or in the absence of Ca^2+^ transporters like the vacuolar Calcium ATPases ACA4/11 or the plasma membrane localized calcium pumps ACA8/10 (Boursiac et al., 2010; Yang et al., 2017). They all display autoimmune phenotypes like stunted growth, activation of defense responses, and constitutive programmed cell death. Interestingly, the *bon1* mutant, which encodes a C2 domain containing copine protein and also displays an autoimmune phenotype, could be rescued by overexpressing hyperactive ACA8 or ACA10 Calcium ATPases that remove Ca^2+^ from the cytosol (Li et al., 2025).

On the other hand, for CNGC2 (DND1) and its closest homolog CNGC4 (DND2), which together form a channel complex (Chin et al., 2013; Tian et al., 2019), the autoimmune phenotype is seen in loss-of-function mutants (Yu et al., 1998; Jurkowski et al., 2004), i.e., in the absence of the channel. These mutants exhibit not only immunity-related phenotypes but also a wide array of defects, ranging from impaired responses to environmental factors like humidity and heat (Finka et al., 2012; Hussain et al., 2024), an attenuated cell damage response (Wang et al., 2022a; Sun et al., 2025), developmental defects like delayed floral transition (Chin et al., 2013), reduced fertility of the sporophyte (Chaiwongsar et al., 2009), and early senescence (Ma et al., 2010) to compromised immune responses (Ali et al., 2007; Tian et al., 2019). This strongly indicates that CNGC2 (and likely its associated CNGC4) plays a broader role beyond immunity.

Interestingly, a double mutant of the vacuolar H^+^ /Cation exchangers CAX1 and CAX3 (*cax 1 cax3*) displays very similar phenotypes to *cngc2/4* mutants, including the stunted phenotype and elevated SA levels (Wang et al., 2017). Both *cngc2/4* and *cax1 cax3* phenotypes are suppressed when the plants are grown in low Ca^2+^ media (0.1mM Ca^2+^, (Wang et al., 2017)), suggesting that Ca^2+^ homeostasis might be the common factor. Indeed, while wild type plants can reset a transient increase in [Ca^2+^]_cyt_ after exposure to external Ca^2+^ concentrations above 10 mM, in *cngc2* and *cax1 cax3* plants [Ca^2+^]_cyt_ levels remain elevated suggesting that Ca^2+^ homeostasis is disrupted in these mutants (Wang et al., 2024). Interestingly, both, *cngc2* and *cax1 cax3* accumulate less total calcium than wild type plants suggesting the issue is not Ca^2+^ uptake but rather distribution or uptake into the cells (Conn et al., 2011; Wang et al., 2017; Wang et al., 2024). It was indeed proposed that *cngc2* and *cax1 cax3* plants are compromised in the unloading of Ca^2+^ from the vasculature into the surrounding cells, leading to an accumulation of Ca^2+^ in the apoplast (Wang et al., 2017). The reason that *cngc2* and *cax1 cax3* plants show the same phenotype could be that Ca^2+^ is taken up from the apoplast into the cytosol by CNGC2 and then further into the vacuole via CAX transporters. If Ca^2+^ cannot be taken up in the absence of CNGC2, Ca^2+^ accumulates in the apoplast (Wang et al., 2017). Similarly, if Ca^2+^ from the cytosol is not moving into the vacuole, the stream of Ca^2+^ is disrupted and again an accumulation of apoplastic Ca^2+^ occurs. In the case of *CNGC2OX* plants, the overexpression of CNGC2 may lead to a constant but slow influx from the apoplast, which does not trigger autoimmune responses but depletes the pool of free Ca^2+^ in the apoplast, which leads to a reduced Ca^2+^ influx upon activation of immune responses, leading to a reduced PTI and increased pathogen susceptibility. The amount of free Ca^2+^ in the apoplast is limited. The majority of Ca^2+^ is bound to biomolecules like pectin or oxalate (Wdowiak et al., 2024).

*cngc2* plants display autoimmune phenotypes, including elevated SA levels and enhanced pathogen resistance to virulent and avirulent pathogens (Yu et al., 1998). At the same time, *cngc2* is deficient in its PTI response, including Ca^2+^ influx upon flg22 treatment (Tian et al., 2019). The *CNGC2OX* lines complemented the *cngc2* autoimmune phenotypes and exhibited a wild type morphological phenotype as well as wild type levels of SA. However, they were more susceptible to virulent pathogens than wild type plants. Unexpectedly, they also displayed a similar attenuated ROS burst and Ca^2+^ influx response upon flg22 treatment as *cngc2*. Further, they were even more susceptible than *cngc2* to elevated external Ca^2+^. This suggests that when CNGC2 levels are above or below a certain level, Ca^2+^ homeostasis is altered leading to aberrant responses. The accumulation of Ca^2+^ in the apoplast of *cngc2* due to a defect in loading it into the cells leads to an attenuated PTI, while the overexpression of CNGC2 may result in too low levels of free Ca^2+^ in the apoplast, and this also leads to an attenuated PTI response.

Previously, it was shown that *cngc2* mutants also display defects in auxin perception and biosynthesis. Most of the *cngc2* phenotypes were partially suppressed in double mutants of *cngc2* with *yuc6*/*rdd1* (*YUCCA6*) or *wei8* (encoding *TAA1*, *TRYPTOPHAN AMINOTRANSFERASE OF ARABIDOPSIS*), which encode the enzymes responsible for the last steps of auxin biosynthesis from tryptophan (Mashiguchi et al., 2011; Chakraborty et al., 2021). Interestingly, our *CNGC2OX* lines also displayed elevated levels of IAA, similarly to *cngc2*. This suggests that a Ca^2+^ imbalance between apoplast and cytosol may affect other cellular processes as well.

In this study, we report that CNGC2 may not be specific for immune responses but rather plays a more universal role—possibly facilitating Ca^2+^ uptake from the environment to support Ca²⁺ homeostasis, which is important for proper cellular signaling. The only alternative hypothesis we can propose at this stage is that CNGC2 and likely its associated unit CNGC4 may function as common subunits utilized across various CNGC heterotetrametric channels, serving as non-specialized support components. We are currently investigating these possibilities further to better understand their biological functions.

## Methods

### Plant Materials and Growth Conditions

For phenotypic analyses, *A. thaliana* seeds were cold stratified at 4°C for 2 days prior to being grown on Sunshine-Mix #1™ (Sun Gro Horticulture Canada). Plants for pathogen infections were grown in a growth chamber with a 9-hour photo period (9 hour light/15 hour dark) and a day/night temperature regime of 22°C/18°C. For Ca^2+^ sensitivity experiments, plants were grown on petri dishes with ½ Murashige and Skoog (MS), 1% sucrose, and 0.8% (w/v) agar at pH 5.8 under ambient light conditions. 20 mM CaCl_2_ plates were made by supplementing ½ MS media with 20 mM CaCl_2_. Plants were grown on CaCl_2_ supplemented plates for 3 weeks and fresh weight of above ground tissue was recorded.

### Generation of transgenic lines

*CNGC2* was cloned into the pEarleyGate 101 plasmid (Earley et al., 2006) to create the *CaMV35S CNGC2* construct. The *CNGC2OX* transgenic plants were created by *Agrobacterium tumefaciens*-mediated transformation of cngc2 plants with the *CaMV35S CNGC2* construct by the floral dip method (Clough and Bent, 1998).

### Pathogen infections

Bacterial infection with *Pseudomonas syringae* pv. *tomato* DC3000 was conducted as reported previously (Yoshioka et al., 2006) using 5- to 6-week-old plants and an inoculum of 5 × 10^4^ CFU ml^−1^. Infection with *Hyaloperonospora arabidopsidis* isolate Noco, which is virulent to Columbia ecotype of Arabidopsis was performed as described previously with 5 × 10^5^ spores ml^−1^. Detached infected leaves were vacuum infiltrated with trypan blue stain (for cell death and pathogen infection) before being boiled for 2 minutes and incubated in dye solution overnight. Chloral hydrate was used to de-stain the samples (Chin et al., 2013).

### Analysis of endogenous Salicylic acid

Endogenous SA was analyzed using the *Acinetobacter* sp. ADPWH lux-based biosensor as described (Defraia et al., 2008).

### Auxin

Levels of endogenous auxin were measured as described (Preston et al., 2009).

### Analysis of flowering transition time

Arabidopsis WT and mutant plants were grown on Sunshine-Mix #1 (Sun Gro Horticulture, Vancouver, Canada) in a growth chamber under a 16-h photoperiod (16-h light/ 8-h dark) at 22^◦^C/18^◦^C. Observations were made every other day and floral transition was recorded as days for the first bolt to form from time of sowing as described in (Fortuna et al., 2015).

### Cytosolic Ca^2+^ measurement

The GCaMP3 expressing line (Toyota et al., 2018) was used for the analysis of [Ca^2+^]_cyt_. Leaf discs of 4mm diameter harvested from 4–5-week-old plants expressing GCaMP3 were floated in individual wells of a black 96-well chimney F-bottom microplate plate (Greiner, 655076) containing 100μL H_2_O. After incubation of 2-3 hours, 100μL of treatment was added to each well. Fluorescence was measured over a period of 1-4 hours using a Tecan Infinite M1000 Pro plate reader using the 488 nm excitation peak and 515nm emission wavelengths, as described previously (DeFalco et al., 2017). Absolute fluorescence values for each well were normalized to the first reading value, shown as F/F_0_ (F represents absolute fluorescence reading and F_0_ reading at timepoint 0).

### ROS analysis

Apoplastic ROS was measured with a chemiluminescence system. Leaf discs of 4mm diameter harvested from 4–5-week-old plants were floated in individual wells of an opaque white 96-well chimney F-bottom microplate (Greiner, 655075) containing 200 μL ddH2O. After 16-18hr incubation in the dark, the water was replaced with 100 μL working solution of 17mg/uL luminol (Sigma, A8511), 0.02mg/mL horseradish peroxidase (Sigma, P6782) and 100nM flg22. Luminescence values were measured with a plate reader (TECAN Infinite M1000 Pro).

### Q-PCR

RNA extraction was carried out using the RNeasy Plant Mini kit (Qiagen) according to the manufacturer’s instructions. Quantitative real-time PCR (RT-qPCR) was performed as described previously (Loranger et al., 2025) using *Glyceraldehyde-3-Phosphate Dehydrogenase C2* (*GAPC2*) and *Elongation factor 1a* (*EF-1a*) gene expression for normalization. Primer sequences are listed in Supplemental Table S2.

## Acknowledgements

This work was supported by a Discovery Grant from National Science and Engineering Research Council (NSERC) to K.Y. and a graduate student fellowship from NSERC to S.C.

